# A divide and conquer metacell algorithm for scalable scRNA-seq analysis

**DOI:** 10.1101/2021.08.08.453314

**Authors:** Oren Ben-Kiki, Akhiad Bercovich, Aviezer Lifshitz, Amos Tanay

## Abstract

Scaling scRNA-seq to profile millions of cells is increasingly feasible. Such data is crucial for the construction of high-resolution maps of transcriptional manifolds. But current analysis strategies, in particular dimensionality reduction and two-phase clustering, offers only limited scaling and sensitivity to define such manifolds. Here we introduce Metacell-2, a recursive divide and conquer algorithm allowing efficient decomposition of scRNA-seq datasets of any size into small and cohesive groups of cells denoted as metacells. We show the algorithm outperforms current solutions in time, memory and quality. Importantly, Metacell-2 also improves outlier cell detection and rare cell type identification, as we exemplify by analysis of human bone marrow cell atlas and mouse embryonic data. Metacell-2 is implemented over the scanpy framework for easy integration in any analysis pipeline.

## INTRODUCTION

Since its initial development over ten years ago, single cell RNA-seq scaled rapidly from the laborious and manual construction of few dozens of Smart-seq^1^ libraries to fully automated and highly parallelized production pipelines^2,3^ capable of generating millions of single cell profiles on diverse applications^4,5^. The characteristics of scRNA-seq profiles remained however largely unchanged since the deployment of unique molecular identifiers (UMIs) for noise reduction^6,7^. Each scRNA-seq profile is characterized by a sparse sample of RNA molecules, where the majority of genes are not sampled at all, or sampled in few copies. The inference of transcriptional programs^8–10^ and dynamics^11–14^ at high quantitative resolution using methods of increasing sophistication^15–19^ relies heavily on the ability to group these sparse profiles together.

We previously introduced Metacell^20^ as a strategy for partitioning scRNA-seq data into disjoint subsets (called *metacells*) that ideally represent repeated sparse sampling from the same multinomial distribution as expected from a recurrent cell state. The rationale underlying the metacell approach is that the summary of transcriptional maps (or manifolds) using metacells, rather than single cells, lowers the risk for smoothing artifacts (compared to imputation approaches), while still maximizing sensitivity and resolution (compared to more coarse-grained clustering). This strategy becomes particularly effective when a large number of cells are sampled. It is thereby important to ensure its scalability, as common scRNA-seq datasets are increasing in size from thousands to millions of cells.

Here we introduce a new and greatly improved Metacell algorithm (MC2) that supports unlimited scaling, using an iterative divide and conquer approach. In addition to the divide and conquer scheme, the algorithm uses a new graph partition score to avoid time-consuming resampling and directly control metacell sizes, implements a new adaptive outlier detection module for better robustness, and employs a rare-gene-module detector ensuring very high sensitivity for detecting transcriptional states that are present in as little as 0.01% of the data. Our efficient implementation of the MC2 algorithm (https://pypi.org/project/metacells/) can quickly compute metacells from any matrix to power quantitative and robust downstream analysis using scanpy^21^, Seurat^22^ or other toolsets^23,24^, or interactive visualization with the metacell viewer/annotator (https://tanaylab.github.io/MCView/).

## RESULTS

### Scalable metacell analysis using two-phase divide and conquer (DAC)

MC2 works in two phases, where in each phase the algorithm is recursive and parallelized (**Fig 1**, details in **Fig S1**). The first phase produces low quality metacells and groups them into metagroups. This is done by a) working independently (and in parallel) across a random partition of cells into small (∼10,000 cells) *piles*, b) screening for outlier cells in all piles and applying the algorithm recursively on (one or more) piles from them to force their grouping into metacells, and c) applying the algorithm recursively to the mean profiles of all derived metacells to generate metagroups. The output of phase I is a set of metacells and grouping of these metacells into metagroups. The second phase recomputes metacells, but uses piles constructed from the metagroups computed by the first phase instead of random piles. Phase II is also invoked recursively on outliers from all piles, but this time channels strong outliers to remain ungrouped. MC2 final output is therefore a set of metacells (from phase II) and identified “final” outliers.

**Figure 1:**
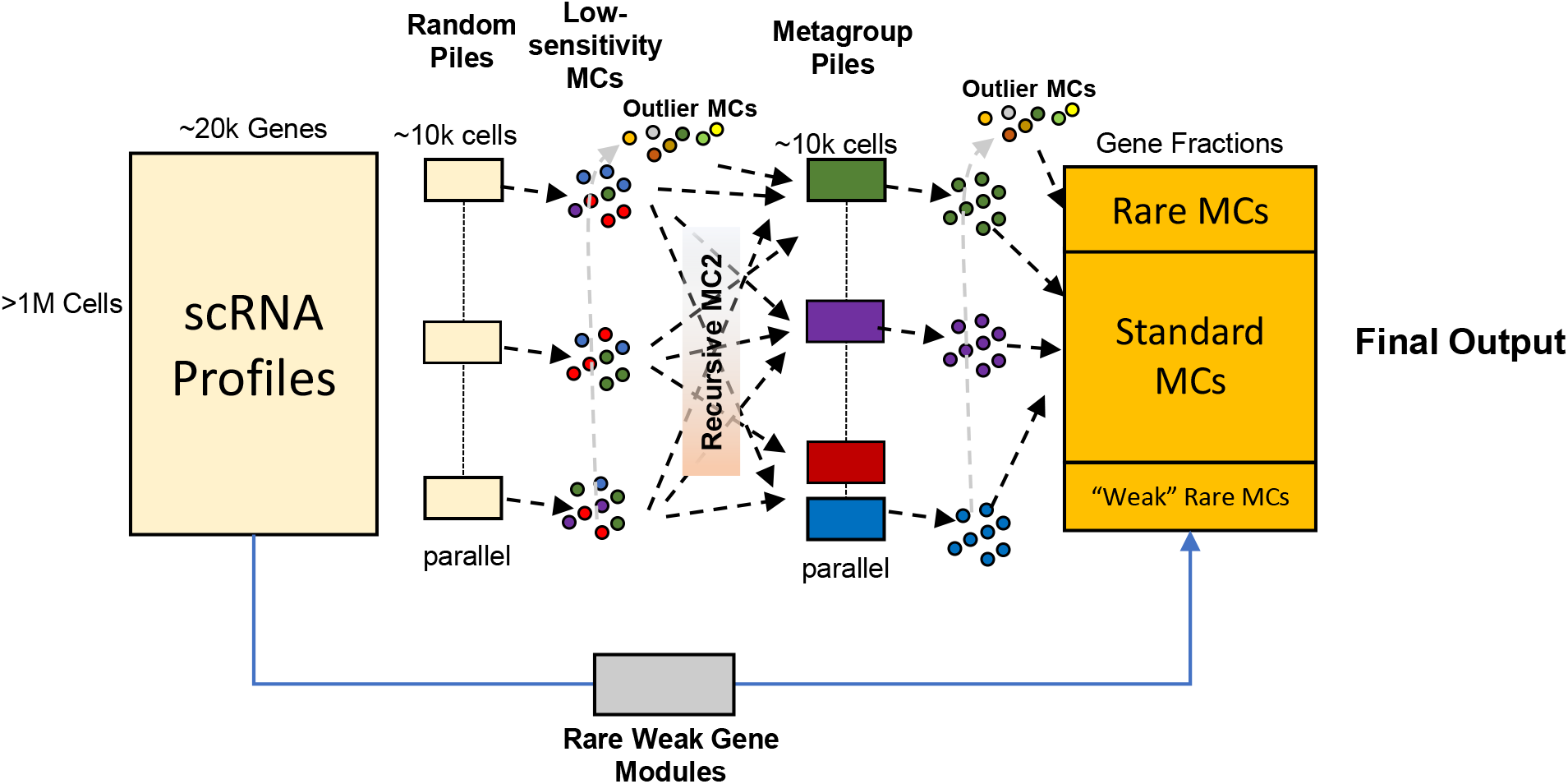
Schematics of the MC2 algorithm. The input is a large UMI matrix and the output is a partition of cell into metacells and final outlier cells. MC2 is deriving a solution using recursive two-phase process. It first divides the data into random piles, and generates low-quality metacells and outliers from them. It then (recursively) groups low-quality metacells into coherent piles, and repartitions these piles to generate high quality metacells. The algorithm is ensuring high sensitivity of rare behavior detection by identifying rare metacell through a pre-process, as well as through regrouping of outlier cells that are being pooled from all piles at both phases of the algorithm.

Scaling of the algorithm is facilitated by working recursively – as long as there are more elements (cells, outliers, metacells, metagroups of metacells) than the amount fitting within one pile, the algorithm will subdivide them, independently (in parallel) group the elements in each pile, collect the resulting groups from all piles, and invoke itself recursively on these groups. This approach allows the algorithm to scale (*O*(*N logN*)) with the number of cells in run time. It also keeps the algorithm space complexity effectively constant (although our implementation uses some compact structures that are linear in the number of cells). Importantly, MC2 naturally enables a high degree of parallelism between piles.

A key consideration in the design of MC2 is the need for sensitivity to detect rare transcriptional states. The MC2 algorithm provides two mechanisms to address this. First, the MC2 outlier detection scheme can trace single cells from rare cell types in random piles, whenever these are not frequent enough for deriving a valid, coherent metacell. The algorithm then pools such bona-fide outliers in outlier piles that are becoming enriched for rare behaviors, leading to their grouping into cohesive metacells when the algorithm is applied recursively on these special piles. This scheme is highly effective as demonstrated below, but can under-perform for rare cell types that are linked with weakly expressed gene markers rather than clear outlier expression profiles. MC2’s second mechanism of rare cell type detection addresses this problem using a gene-based strategy. It runs a pre-process that screens for gene modules with weak but significant correlation structure over all cells, identifies cells expressing specifically such modules, and forms metacells from them prior to the application of the full MC2 two-phase procedure.

In summary, MC2 avoids PCA, global K-nn graph derivation, or the construction of quadratic scRNA-seq similarity matrices, and instead breaks the metacell derivation problem into smaller problems that are being refined as the algorithm identifies which cells should be analyzed together. The algorithm sensitivity relies on a combination of explicit rare behavior detection and a hierarchical method for filtering and grouping rare cellular states.

### MC2 is sensitive and robust

We tested the robustness and sensitivity of the MC2 algorithm in a series of comparisons. Idealized metacells represent cells sampled from the same multinomial distribution and should therefore have intrinsic gene variance proportional to the gene mean. We therefore assessed quality of metacell solutions by quantifying the degree of normalized variance per gene (*inner normalized variance*).

We first wished to ensure that the MC2 algorithm’s efficient graph partition algorithm is not losing significant quality compared to the original, resampling-based Metacell implementation (MC1) ^20^. MC2’s graph partition is applied within each pile, and provides tight control over metacell sizes (which is measured in the total number of UMIs in addition to the number of cells). On the other hand, MC1’s usage of resampling iterations, while not scaling well to large datasets, can potentially enhance robustness. For direct comparison, we applied MC2 in one pile (no divide and conquer) to 160K peripheral blood single cell profiles on which MC1 was applied before. We observed comparable inner-normalized-variance in the MC2 direct partitioning algorithm compared to the MC1’s resampling version (**Fig 2A, Fig S2**). Somewhat lower variance was derived when using the divide and conquer mode of MC2 compared to the single-pile version (**Fig 2B**). MC2 2D visualization of large-scale data is based on plotting metacells rather than cells, facilitating ease of interpretation (**Fig 2C**). These data confirm MC2 and in particular the divide and conquer strategy is deriving metacells at quality that is comparable to the original MC1 implementation for small datasets, while allowing unlimited scaling and better control as discussed further below.

**Figure 2:**
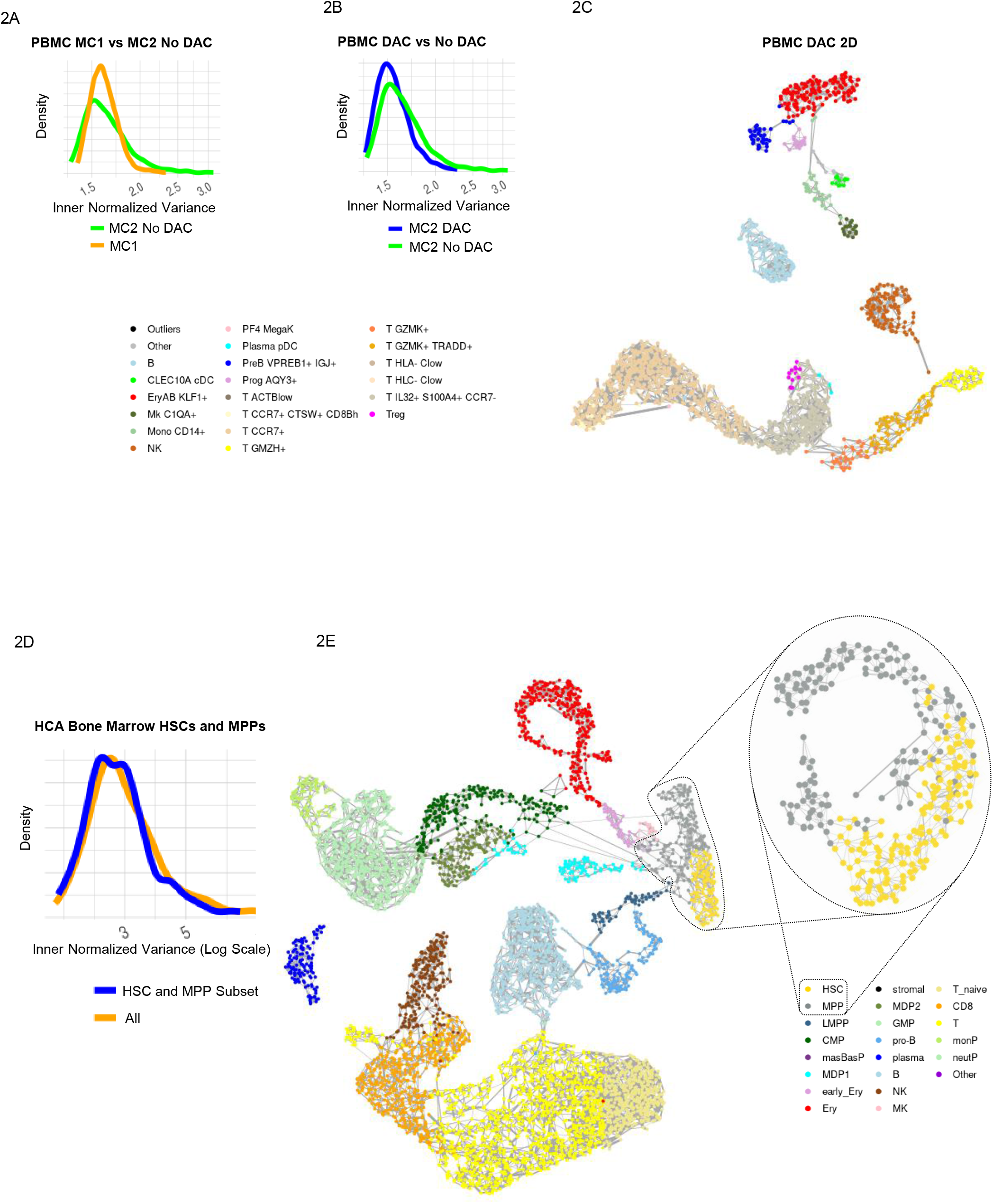
Robustness of the divide and conquer metacell algorithm. A. Distribution of metacell normalized inner variance for the PBMC dataset, using the Baran et al. algorithm (orange) vs. MC2 two-sided stability score optimization, working on the entire data in a single pile (i.e. no divide and conquer, green). B. Distribution of normalized inner variance for the PMBC dataset using the full MC2 algorithms (blue) vs. the single pile algorithm (green). C. Metacell graph derived by MC2 on the PMBC dataset. Annotation as in Baran et al. D. Distribution of metacell normalized inner variance for HSC and MPP cells, when using full MC2 on the HCA bone marrow data set (orange) or when restricting analysis to MPP/HSC cells alone (blue). E. Metacell graph for the full HCA BM data set, and for metacells computed on the zoomed-in HSC/MPP subset.

To test how well MC2 maintains local accuracy when working on a large dataset, we studied ∼380K human bone marrow cells from the human cell atlas^25^. We first applied MC2 to the entire data. We then identified in a supervised fashion all metacells with HSC or MPP characteristics (**Fig S3A-B**), and generated a smaller dataset including the 6666 cells from these metacells. We then compared the metacells derived by MC2 to a set derived by applying the algorithm on a single pile consisting only of HSC/MPP cells (**Fig 2D-E**). This confirmed metacell cohesiveness is maintained when analyzing large datasets, demonstrating the MC2 approach is not losing sensitivity compared by the direct (but less scalable) approach.

### MC2 is scalable to millions of scRNA profiles

We next tested the scalability of MC2 on datasets with millions of cells. We applied MC2 to ∼1.8M cells acquired from mouse embryos during organogenesis stages^26^ (E9.5-E13.5), and compared the results to current popular pipeline using PCA and two rounds of Louvain clustering as implemented in Seurat^7^. Using a single workstation with dual CPUs of 14 cores each, we observed MC2 provides a major reduction in elapsed time (∼40 minutes for MC2, vs. ∼150 minutes for PCA + 2-level Louvain clustering on ∼1.8M cells). The algorithm also supports graceful scaling in maximal memory (**Fig 3A-B**) that is permissive for running MC2 on much larger datasets. Much of the improvement in elapsed time is due to better parallelism of the algorithm, showing potential for further (essentially unlimited) scaling when using more than one compute node.

**Figure 3:**
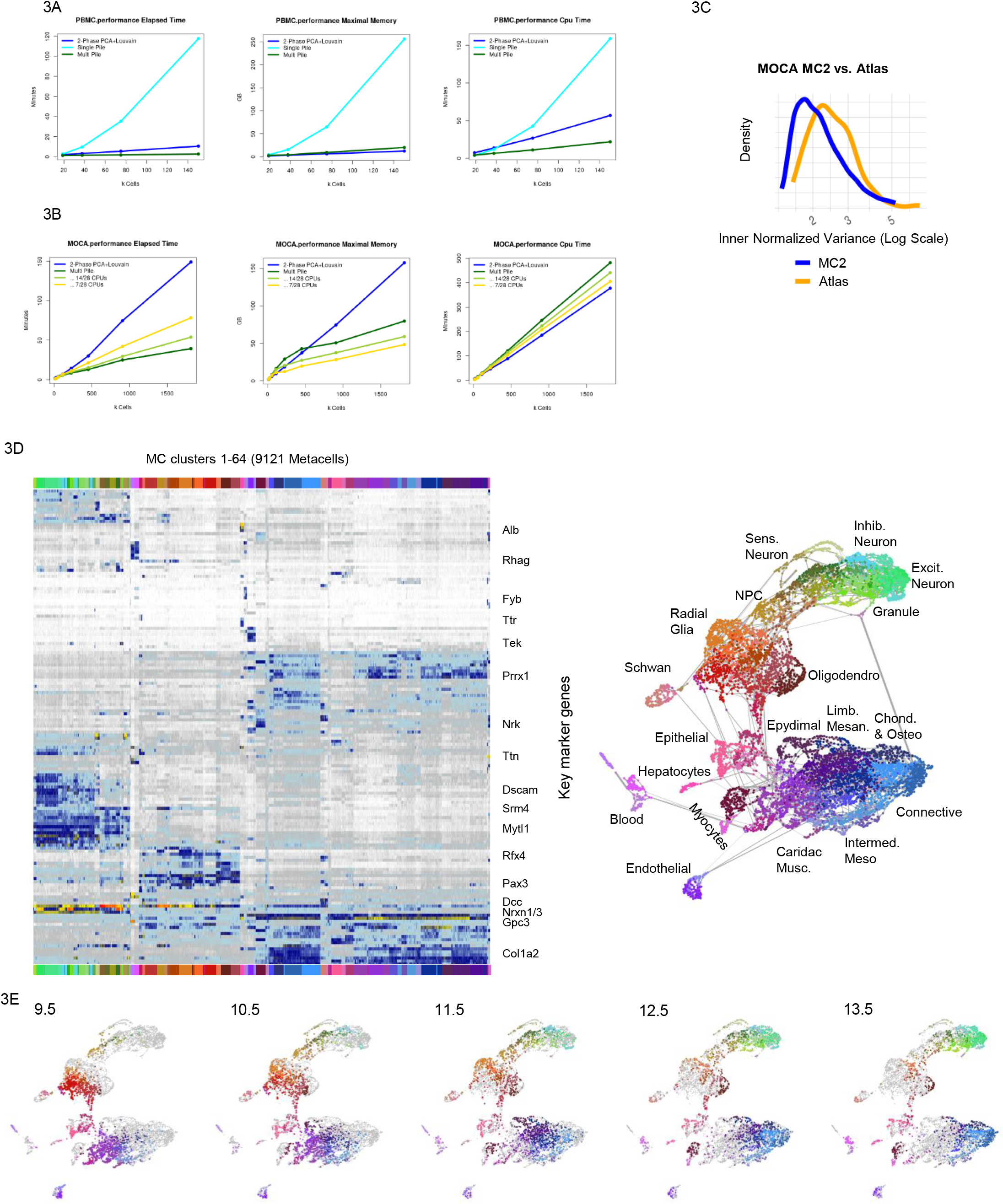
Scaling MC2 to millions of cells. A. Graphs show scaling of MC2 (multi-pile) compared to a naïve metacell on a single pile or a PCA+2-Phase Louvain clustering implementation in Seurat, using the PMBC 160K cell data (resampled to datasets of increasing sizes – X axis). B. Comparison of MC2 and two-phase clustering performance for the organogenesis datasets (MOCA). C. Distribution of normalized inner variance for MC2 and PCA+Louvain original sub-clusters on the organogenesis data. D. Marker heat map and metacell graph projection of the organogenesis data. Clustering of metacells is used for coloring and cross-reference purpose, in support of, but not in place of supervised annotation. E. Distribution of metacells linkage with different embryonic time points over the metacell graph. Color coding is based on metacell clustering as in D. To compensate for differences in the number of cells we randomly sampled 2000 points for each time point, weighted by the fraction of the cells of each age in each metacells.

While it is natural to compare scaling of MC2 to the scaling of two-stage PCA+Louvain clustering, the output of MC2 is different from the clustering solutions. It provides a high granularity model that is designed for use in downstream (quantitative) analysis and not as a substitute for cell typing and sub-typing. In particular, two-phase clustering of the organogenesis dataset is not fully eliminating high variance within the 693 sub-clusters defined, compared to the higher granularity and more precise estimation of quantitative states facilitated by the 9121 inferred metacells on the same data (**Fig 3C**). Following iterative elimination of doublet cells (Methods), MC2 provided a metacell cover that supports a highly informative visualization of the organogenesis manifold (**Fig 3D**). Clustering of metacells using their parametric gene expression state approximated the transcriptional space using 64 large scale behaviors (K-means clusters), which were generally but not perfectly consistent with previous cell type annotation (**Fig S4)**. The derived model reflected temporal dynamics (**Fig 3E**) and broad germ-layer structure that is missing from common single cell PCA+UMAP visualization. The derived structure also facilitates high-resolution in-depth characterization of the combinatorial and quantitative transcriptional gradients within types, as exemplified for epithelial or endothelial cells (**Fig S5**). Further analysis of such large scale metacell models is ideally interactive, as facilitated by our MCView web-based visualizer program (https://tanaylab.github.io/MCView/). Overall MC2 efficiently converts very large scale scRNA-seq data into building blocks that can be used to create a working model of the underlying transcriptional manifold.

### MC2 outliers and rare type detection

One of the major challenges in analyzing very large scale scRNA-seq is the sensitive detection of rare behaviors. Such behaviors may be lost when sub-sampling data, and can require more statistical power for detection within vast samples of less informative recurrent states. MC2 uses two mechanisms for detecting rare behaviors, the first involving a pre-process that searches for rare gene modules, and the second using the MC2 divide and conquer algorithm outlier detection and the recursive analysis of detected outliers for regrouping and inference of rare metacells. We screened the organogenesis metacell cover to identify genes expressed in one metacell at least 8-fold higher than in 99.8% of all other metacells. This resulted in identifying 260 genes spanning over 30 clusters with two or more genes, each representing a defined rare cell state (**Fig 4A**). 47 of these genes were detected during the pre-process stage of MC2, while all others were detected within piles of common outliers or as specific metacell state within a coherent pile.

**Figure 4:**
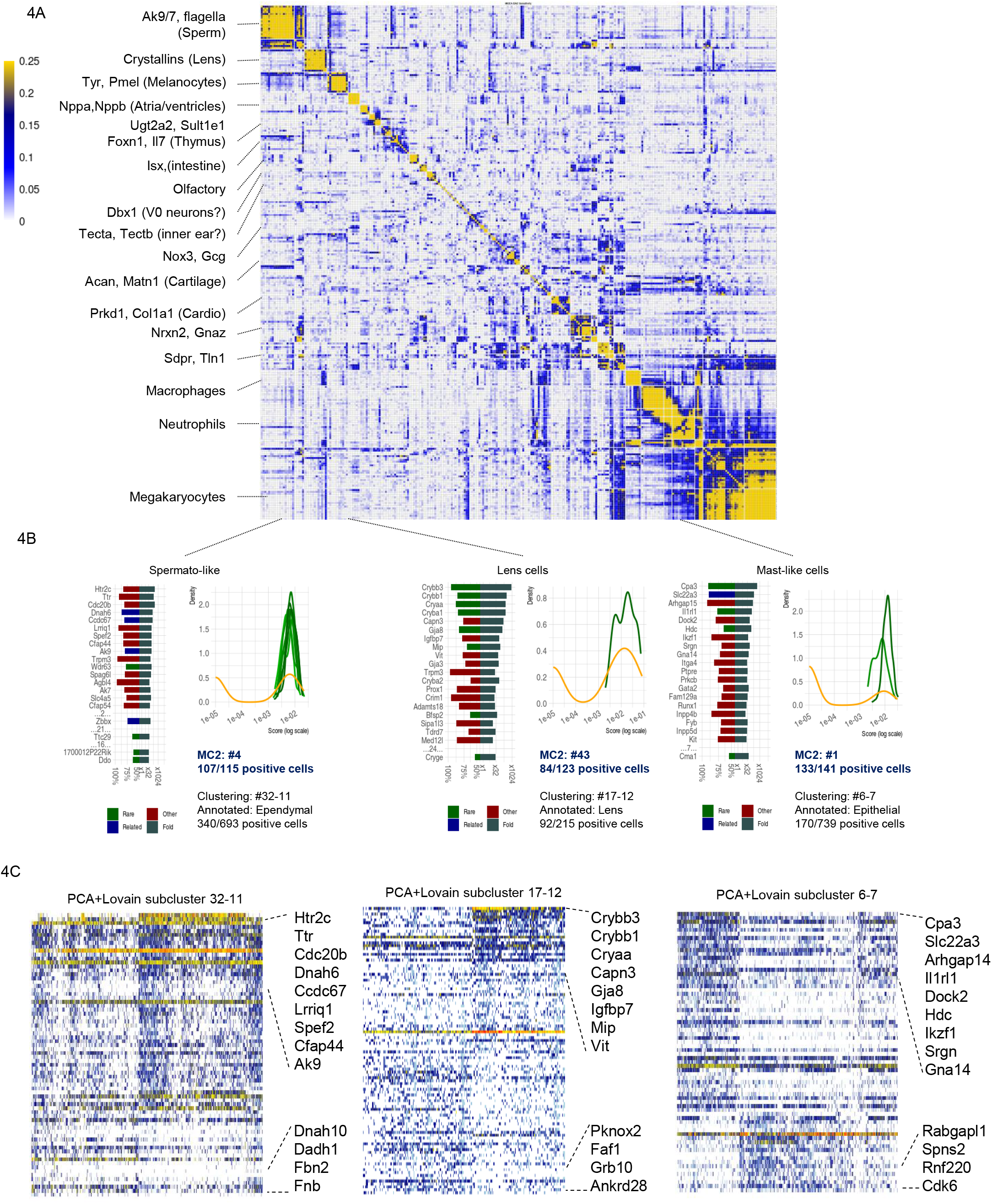
MC2 sensitive detection of rare behavior. A. Correlation matrix between log gene expression frequency of 260 genes with rare expression signatures (see text). We highlight several gene clusters at the left. B. Each bar graph show specificity (left) and fold change enrichment (right) of top genes separating three exemplified rare transcriptional behaviors. Also shown for each rare behavior are the distribution of total gene expression of rare genes per single cells within the top-enriched metacell(s) (shades of green) and within the top enriched PCA+Louvain subcluster (orange). C. Shown are single cell gene expression for rare behavior marker genes and for genes correlated and anticorrelated with them, plotted for cells within the most strongly enriched PCA+Louvain sub-cluster for the observed behavior.

**Figure 4B** demonstrates MC2’s sensitivity and specificity on three rare behaviors involving Lens cells, Mast cells and Spermatogenic-like cells, all of which were identified through rare gene modules pre-screening. MC2 first identifies genes with globally very low mean expression but still significant correlation to other genes over a few cells. Groups of such correlated genes are used to form *rare gene modules*. MC2 then searches for additional genes that are enriched in cells expressing the module, and then moves all cells with potentially significant expression of any (expanded) rare gene module to a specialized pile in which rare metacells (and outliers) are being inferred. The accuracy of detecting rare behaviors can be substantially higher than the accuracy enabled by the standard two-phase clustering approach. In PCA + clustering, some of these rare behaviors are absorbed within subclusters that mix specific rare cells with other, non-specific cells (**Fig 4C**). In conclusion, using a combination of a sensitive pre-process and the divide and conquer strategy, MC2 ensures high sensitivity for detecting rare transcriptional states while scaling naturally to very large datasets.

## DISCUSSION

We have introduced a new scalable algorithm for inferring metacell covers on large scRNA-seq data, demonstrating its robustness and sensitivity for analysis of challenging datasets involving millions of cells. Metacells are groups of single cell profiles that provide sufficient coverage for inference of one quantitative transcriptional distribution. Ideally, profiles within a metacell should be distributed as if the only source of variance in the data is the sparsity of RNA sampling from single cells. As datasets becomes larger and sampling of transcriptional states becomes saturated, this assumption becomes progressively more realistic. Larger data thereby provide strong justification for analysis of metacell transcriptional states (which are quantitative and of more convenient size) rather than direct analysis of single cell profiles and their K-nn graphs.

The implementation of MC2 and the original metacell algorithm is tuned for the typical distributions observed in scRNA-seq, and their application to other single cell genomics data (e.g., scATAC-seq, scBIS-seq) is recommended only if adequate similarity metrics and feature selection strategies are developed. Such adaptation is not described here. It is however natural to use the divide and conquer strategy introduced here for scaling analysis of large-scale single cell omics of multiple types.

MC2 provides effective building blocks for understanding complex transcriptional manifolds. Metacells’ transcriptional states can be assumed to be quantitative and describe the distribution of gene expression in an idealized cell state given sufficient sampling and guaranteed homogeneity. The MC2 underlying model remains however non-parametric and extremely simple, as it avoids any assumptions on the linkage between metacell states. Further work must be channeled toward refined models of the linkage between transcriptional states, but such work, in our mind, should move away from the K-nn, non-parametric approaches that are commonly used in the literature, and toward a principled and quantitative model that put transcriptional states and the connection between them in an interpretable (and ideally mechanistic) context. With more parametric models linking metacells into proper quantitative models, it will be possible to envision the routine usage of large-scale transcriptional atlases as universal references for the interpterion of experiments generating new scRNA-seq data following perturbation, stimulation, patient sampling and more.

## METHODS

### Rare Gene Modules Detection

MC2 primarily detects rare cell types by screening through random data partitions while classifying and grouping outlier behaviors as described below. But in some cases, low UMI count in marker genes of rare behaviors makes it impossible to detect rare cells as outliers in random subsets. To handle such cases, MC2 implements a rare gene module detector that efficiently pre-processes the entire UMI matrix and enhances sparse gene features. This stage detects rare gene modules, collects the cells that express these modules, and invokes the divide-and-conquer algorithm on the cells of each such module separately from the rest of the cell population. The resulting metacells are passed to the final output. The overall flow is working as follows (default main parameters are also specified in **Table S1**):

1. Identify *rare genes* – genes that are observed in a small fraction of the cells (default p=1e-3), but are still observed abundantly (at least 7 UMIs) in at least one cell.
2. We compute the correlation matrix of between all rare gene expression *r*, and then compute its second order correlation matrix defined as *r*_2_ = *cor*(*r*). We then perform hierarchical clustering of *r*_2_ using Ward’s method.
3. For each subtree of the hierarchical clustering we compute the mean *r*_2_ of gene pairs within it. We next consider all maximal sub-trees on at least 4 genes with mean *r*_2_ of at least 0.1 as candidate gene modules. We next repeat the following stages (4-7) for each candidate gene module M.
4. Identify all cells C with one or more UMIs from the genes in M.
5. Add to M all genes whose UMI frequencies in the cells C are at least 128-fold higher than their frequencies over all cells, as long as this increases the number of cells expressing the gene module by a factor of less than 4.
6. Screen for all cells (including C and others) with at least 4 UMIs observed for genes in the (expanded) module M. Denote these as *C*^*M*^.
7. Modules for which|*C*^*M*^| < 12 are discarded since these do not suffice to create even a single metacell. Modules for which |*C*^*M*^| > *T* are considered to be too common and are discarded as well. T is defined as the total number of cells required to give rise to (on average) at least 48 cells in a random pile (as described below).

All the threshold parameters used above are tuned to maximize complementarity between this rare gene module detection pre-processing, and phases of outlier cell detection and re-clustering within the main MC2 divide and conquer algorithm. We do not anticipate scenarios requiring adjustment of these parameters.

### A two-sided stability score for graph partitioning

MC2 runs by applying graph partition on metacell graphs constructed over piles of cells in the data, or recursively over groups of metacells. The goal is to partition the graph into subsets with high connectivity and homogeneous size distribution. Compared to our previous version of Metacell (MC1), we wish to avoid computationally expensive resampling iterations, and define an explicit score function to stabilize the original local optimization steps and cooling strategy. On the other hand, in contrast to the popular modularity metric^27–30^ and its different flavors, in MC2 we wish to discourage inclusion of a node in a partition if its internal connectivity is very asymmetric (e.g., only outgoing edges to members of the metacell).

Define a cell graph *G* with nodes (cells) V and weighted edges *E, w*_*e*_: *E* → *R*^+^. This graph is constructed as in MC1, using a balanced K-nn graph construction. We will denote the incoming and outgoing neighbors sets as *N*^*in*^(*v*), *N*^*out*^(*v*), and score a partition into M metacells *mc*(*v*): *V* → [0, *M* − 1], *M*_*m*_ = *mc*^−1^(*m*).

We first compute for each node its probability for staying inside a partition assuming a random walk starting from the node (using outgoing edges) or a similar probability assuming the process is working in reverse (using incoming edges):

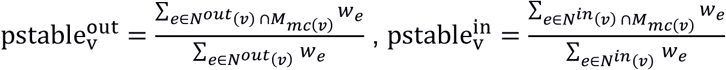

We wish to compare these probabilities to the uniform distribution (assuming a random walk on a fully connected, weight-less graph):

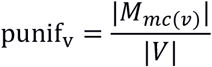

Importantly, we now define two ratios of stability (forward and reverse) separately:

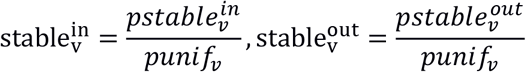

And consider our goal function, denoted as the *two-sided stability score*, by a non-linear (geometric mean) summation of these scores over all nodes:

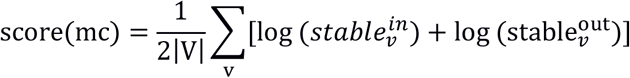

We note that when using this scoring scheme, the cost of keeping an edge inside or outside a partition is not constant as it would be in the modularity metric, which is making optimization somewhat more computationally difficult. Nevertheless, the two-sided stability score does more strongly discourage the inclusion of nodes in partition if their connectivity to the partition is highly non-symmetric.

### Generation of partitions with optimized two-sided stability score

Given a weighted graph *G* = (*V, E, w*), our algorithm is searching for a partition *mc* with an optimized two-sided stability *score*(*mc*) and the metacell size restrictions 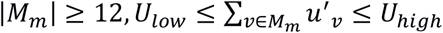. Metacell size is defined by the total number of UMIs in its cells *u*_*v*_, but in order to avoid highly asymmetric cell sizes leading to metacells with very few cells, we cap all *u*′_*v*_ = min (*u*_*v*_, *median*_*v*_(*u*_*v*_) * 2). Restriction on metacell sizes is determined using a user parameter defining the target metacell size *U*_*targ*_, as 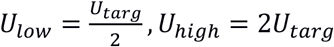. We are however increasing *U*_*targ*_ beyond user request for datasets with large cells, if it implies less than twelve cells on average per metacell. The algorithm works using the following steps:

1. *Seeding*: Choosing 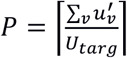 seeds similarly to the MC1 algorithm. This involves iteratively sampling nodes that are disconnected from the nodes selected so far and their neighbors. Seeding ends up with a partition *mc*.
2. *Optimization*: This step is incrementally improving the score by moving nodes between the partitions until no such steps are possible. To improve the optimizer robustness, we start the optimization sequence allowing also sub-optimal changes in node *v* metacell association, by adding to the difference in overall *score*(*mc*_*new*_) additional contribution from difference in the *v* individual stability 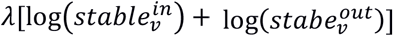. The parameter *λ* is starting at high levels, which are allowing nodes to switch to a partition that provides more connectivity even if this result in reduction in the stability of other nodes in its current partition. The parameter is gradually reduced to 0, and in its final stages the algorithm is directly optimizing the goal function.
3. *Max-Size control*. We identify all metacells exceeding size restriction 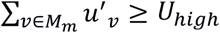 and dissolve each of them. We return to steps 1-2 by re-seeding only the dissolved cells, and re-optimizing the partition (keeping the size-valid metacells initially intact). For efficiency we use *λ* = 0 for all the non-dissolved nodes during this re-optimization. We iterate this until all metacells meet the maximal size restriction.
4. Min-Size control: we identify all metacells with at least one gene showing mean expression that is 8 fold higher than the mean over all metacell. We dissolve all small metacells 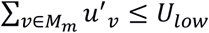 (for metacell without a strong maker gene) or 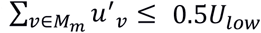 (for metacell with a strong marker gene). In any case we dissolve metacells with |*M*_*m*_| < 12. We then return to steps 1-2 by reseeding them with one less seed than the number of dissolved metacells, and re-optimizing similarly to step 3. We iterate this until all metacells meet the minimal size restriction, or until the next iteration causes the creation of a too-large metacell.

### Improved Deviant (Outlier) Cell Detection

We generally wish to ensure metacells include cells for which all genes are following one multinomial UMI distribution. Previously, we suggested to identify deviant (outlier) cells as those with at least one gene that is severely over expressed (fold factor over 8) compared to the mean expression in the metacell. However, using this criterion can result in massive (or even complete) dissolution of metacells in many datasets, due to high noise level, inter-individual differences, or other effects. We therefore developed a new adaptive deviant cell removal algorithm that tunes two critical parameters: 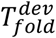 the minimal fold factor of deviant gene expression, and 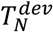 the maximal number of cells that can be defined as deviants based on the expression of one gene.

Given these parameters, we define deviant genes as those with maximal fold factor at least 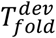, and score each cell using the minimal rank of its fold factors over all deviant genes. Following this, deviant cells are specified as those having minimal rank of at most 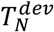. It is easy to see this ensure that no more than 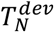 cells are removed due to outlier behavior of any single gene.

To select 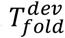, we set its baseline at 8 and increase it while not more than 3% of the genes are defined as deviant for one or more cells.

To tune 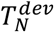 we set its baseline at 1 and increase it while not more than 25% of the cells are deviant.

After removing the deviant cells, we may end with some metacells that are too small. We completely dissolve any such too-small metacells, using the same method as described in step 4 (min-size control) above. We mark all the deviant and dissolved cells as outliers, and move them to later re-analysis within the hierarchical divide and conquer scheme we next describe.

### The Metacell-2 overall algorithm: Divide and Conquer

A divide-and-conquer algorithm is needed to compute metacells from large dataset using a reasonable amount of CPU and memory. The complexity of the direct graph partition algorithm is O(N^2^) since it requires computing similarities between all the cells. The complexity of the divide-and-conquer algorithm we describe below is O(N log N), where N is the number of cells. The algorithm works in three phases (see **Fig 1**, details in **Fig S1**):

#### Preliminary phase

In this phase, we assign the cells to random piles of a manageable size (∼10K cells), and invoke the graph partition algorithm to group them into metacells, followed by detection of outlier cells. We combine the outliers reported by all the piles and repeat the process (random pile-partition and metacell derivation) until we have a final single outlier pile. We group the remaining single outlier pile into metacells while forcing 100% coverage (that is, without removal of outliers). This results in a complete partition of the cells into metacells. However, the quality of this partition, being based on random piles, can be low.

#### Metagroup phase

In this phase, we consider each of the metacells computed by the preliminary phase to be a single observation (using the total UMI counts over each metacell). We recursively invoke the full divide-and-conquer algorithm to group these metacells into metagroups (forcing 100% coverage as above). We note that more than one recursive iteration is needed when the dataset is larger than 0.5M cells (over ∼10K metacells of ∼50 cells each). Also note that in terms of enforcing partition size, at the metagroup phase we restrict partition size between 5,000 and 12,500 cells, rather than *U*_*targ*_ UMIs in the metacell phase.

#### Final phase

In this phase, we construct from each metagroup a new pile (consisting of the cells within its metacells). We now invoke the direct algorithm to compute metacells for each of these piles. We again combine the outliers from all the piles and recursively repeat the process, however in this phase we stop recursion after one level, and designate the outliers detected in outlier piles as *final outliers*.

This concludes the algorithm, which is reporting metacells derived from the rare gene modules pre-process, combined with metacells derived by the main algorithm, and a limited number of remaining un-clustered final outlier cells.

### scRNA-seq data sources and pre-processing

- **PBMC**: We used PBMC160k data as previously described (Baran et al). We excluded all-zero genes, mitochondrial genes, as well as IGHMBP2, IGLL1, IGLL5, IGLON5, NEAT1, TMSB10 and TMSB4X. We then excluded all cells with less than 800 UMIs, more than 8,000 UMIs, or with more than 10% of their UMIs from the excluded genes. This left us with 22,617 out of the original 32,738 genes and 149,825 out of the original 163,234 cells.
- **HCA**.**BM**: We downloaded the HCA bone marrow data from (https://data.humancellatlas.org/explore/projects/cc95ff89-2e68-4a08-a234-480eca21ce79?catalog=dcp1). We excluded all-zero genes, mitochondrial genes, as well as MALAT1 and XIST. We then excluded all cells with less than 800 UMIs, more than 25,000 UMIs, or with more than 30% of their UMIs from the excluded genes. This left us with 27,261 out of the original 33,694 genes and 302,766 out of the original 378,000 cells.
- **MOCA**: We downloaded the MOCA organogenesis dataset^31^. We excluded all-zero genes, mitochondrial genes, as well as MALAT1 and NEAT1. We also excluded 1700007G11Rik, 1700019B21Rik, Cmtm8, Col4a4, Fem1b, Gm11375, Gm28826, Gm43298, Kyat3, Lancl2, Minpp1, Olfr1062, Parn, Poldip3, Sirpb1b, Syt16 and Vmn2r-ps49 as genes which were both “noisy” (had normalized variance/mean above 2.5) and “lonely” (had correlation of less than 0.1 with all other genes). We then excluded all cells with less than 300 UMIs, more than 3000 UMIs, or more than 20% of their UMIs from the excluded genes. This left us with 26,143 out of the original 26,183 genes and 1,798,929 out of the original 2,058,652 cells. We then run the MC2 algorithm, and marked as doublets all cells detected as such in the atlas, as well as all cells inside metacells where at least 1/8^th^ of the cells were detected as doublets in the dataset. This left us with 1,658,637 cells.

All further analysis was done on the above filtered data.

### Metacell QC metrics

An ideal metacell contains UMI vectors that are generated by sampling from the same multinomial distribution. We test how well a given metacell solution is following this hypothesis by computing the normalized variance (variance over mean) over the (downsampled) cells of each metacell, for genes with at least 40 UMIs in total, and take the 95^th^ percentile (over all genes) of these values as the inner normalized variance of each metacell.

The above measure is sensitive to the sizes of metacells and this can skew the results when comparing datasets with different metacells’ size distributions. To mitigate this effect, we adjust metacell size distributions prior to comparison, working separately on each annotated cell type. For each such type, we separately sort the metacells by their size in each dataset. We then adjust metacell sizes (by sampling cells from them) such that the size distributions of metacells in the two datasets is similar. We note that we can robustly compare more than two datasets when this normalization scheme is applied to all of them simultaneously.

### Rare Behaviors

MC2 output includes a set of rare gene modules that are being used to identify rare cell types and derive metacells enriched for them.

#### Distinct Genes

For each rare gene module, we consider as “real positive” the cells that belong to metacells that were computed from cells identified as expressing the module by the MC2 algorithm as described above. We then compute the AUROC for using each gene as a classifier for these cells, and show the highest AUROC and fold factors genes, as well as the original genes, in the module identified by the MC2 algorithm.

#### Score distributions

For each such gene module, we score all cells using the total fraction of UMIs from this module out of all UMIs. Given any cell partitioning (metacells, or sub-clusters), we identify the metacell/subcluster with highest mean score and set a score threshold to be 50% of the median score for cells within it. All cells with scores of at least the threshold are considered “real positive” cases. We next show the distribution of these scores in the metacells/clusters most enriched for these cells.

## Acknowledgments

We thank people in the Tanay group for discussion. Research was supported in part by the ERC (project scAssembly), The EU Braintime project, Chan Zuckerberg Initiative, and the Kahn foundation.

## Figure Legends

**Figure S1:**
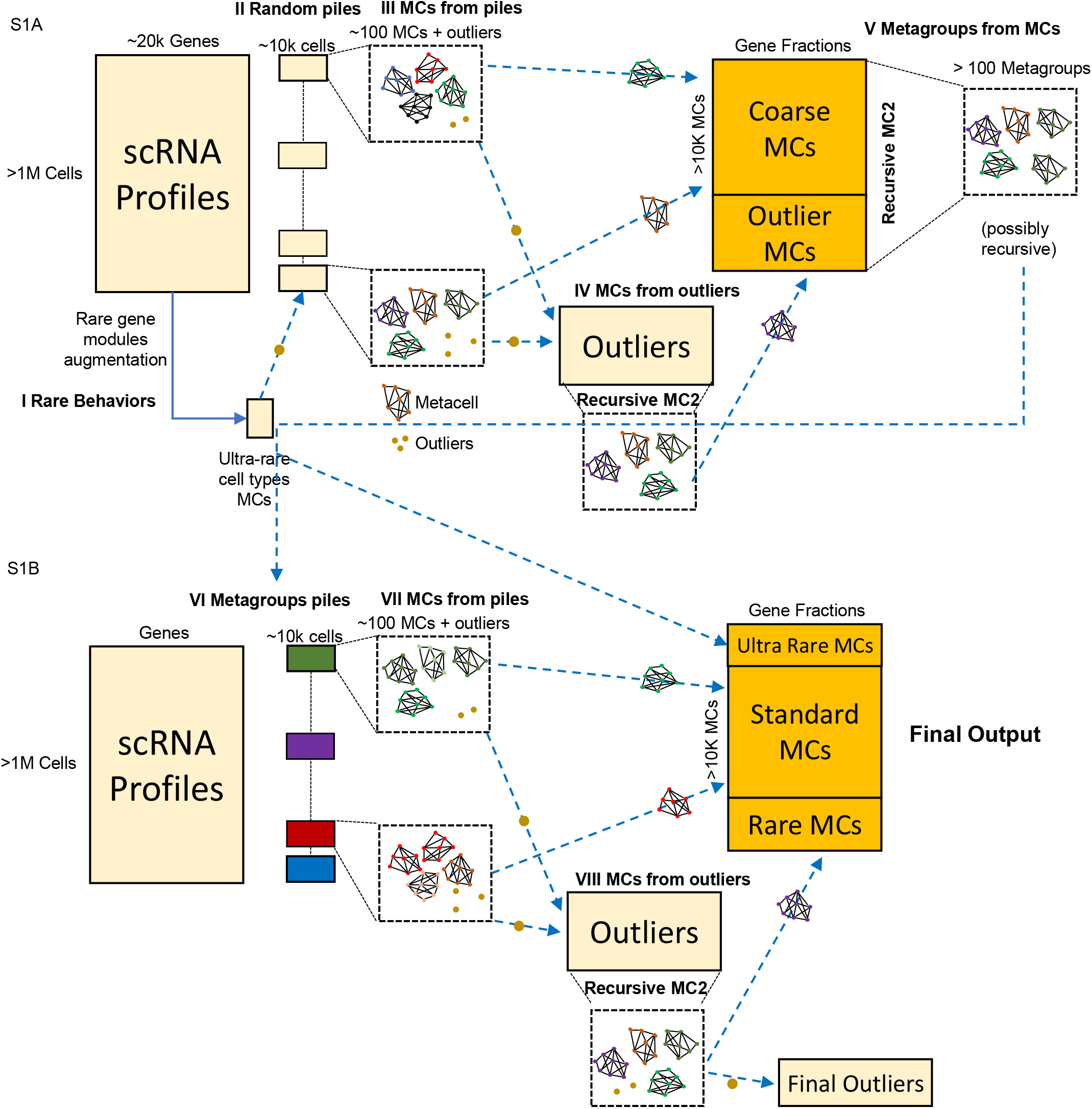
More detailed schematics of the MC2 algorithm. The input is a large UMI matrix, MC2 is a recursive two-phase process working as follows: **I**, Preprocessing detecting rare gene modules and metacells based on them and move these directly to the output, **II**, dividing all cells into random piles. **III** – graph partition defines metacells in each pile. **IV**, outlier cells are removed from metacells into specialized piles, used to create additional (rare) metacells (recursively). **V**, MC2 groups metacells into metagroups (recursively). **VI**, metagroups are used as new homogeneous piles. **VII**, Generation of metacells in 2^nd^ iteration piles. **VIII**. Remaining outliers collected from 2^nd^ iteration piles, further grouped into rare metacells, with detection of final outliers. The final output is a set of final metacells and outliers.

**Figure S2:**
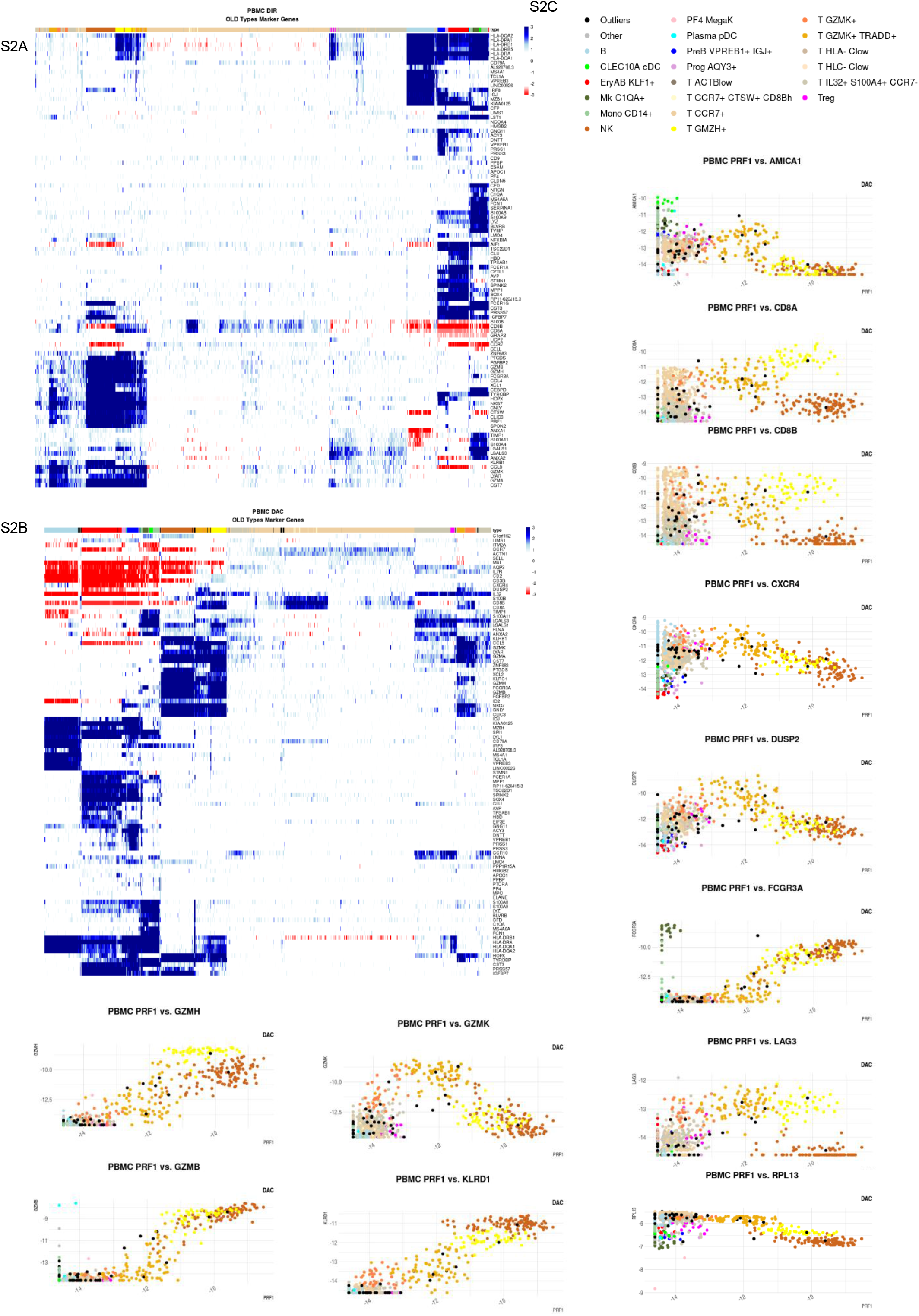
MC2 models for PBMC with and without divide and conquer. A. Heatmap showing marker gene expression (log2 normalized compared to median) for the non DAC MC2 model. B. Heatmap showing marker gene expression (log2 normalized compared to median) for the DAC MC2 model. C. Gene expression per metacell for select T-cell genes as in Baran et al.

**Figure S3:**
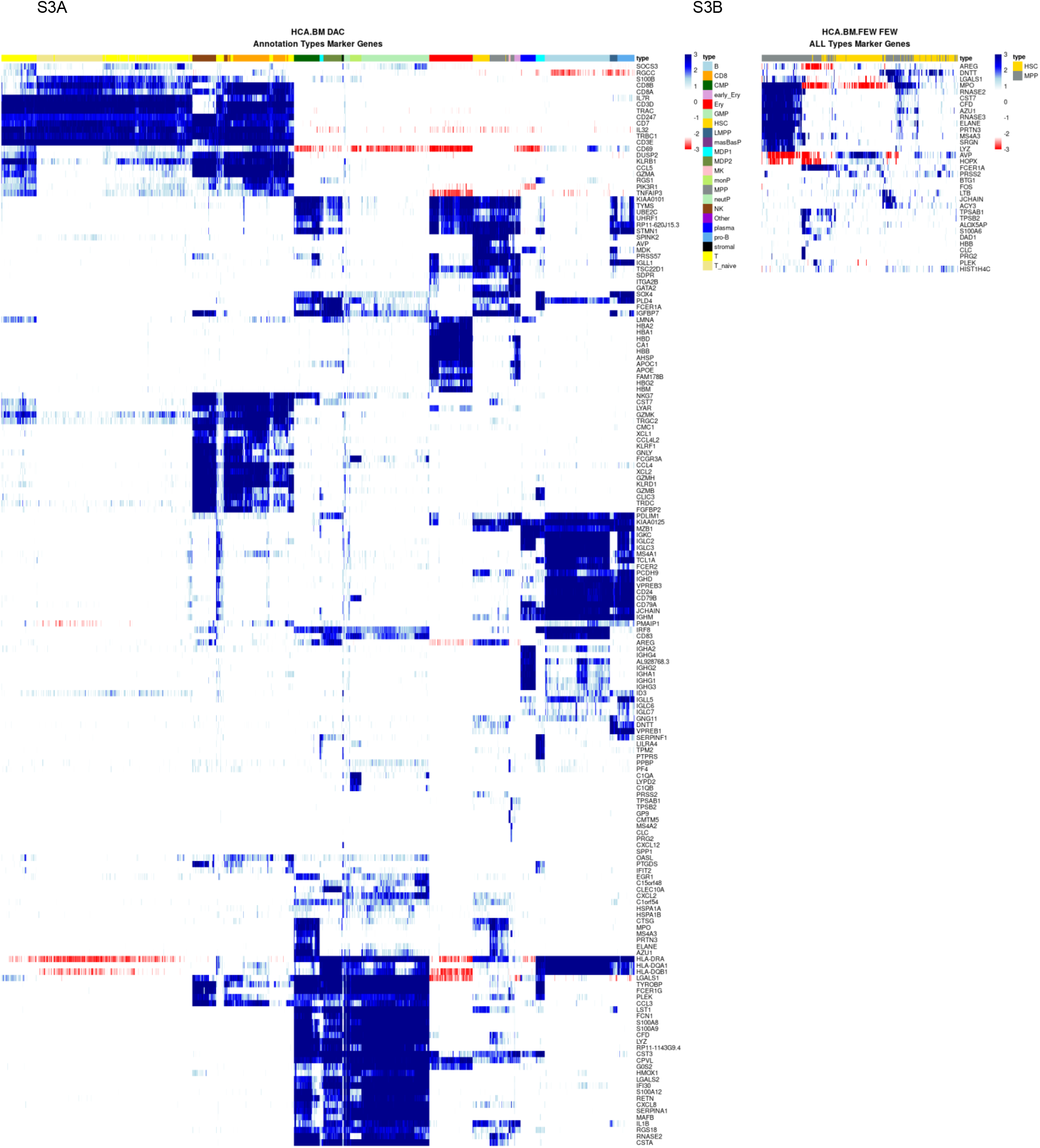
Bone marrow MC models. A. Heatmap showing marker gene expression (log2 normalized compared to median) for the bone marrow global model. B. Marker gene expression for the MC2 models constructed using only HSC/MPP cells.

**Figure S4:**
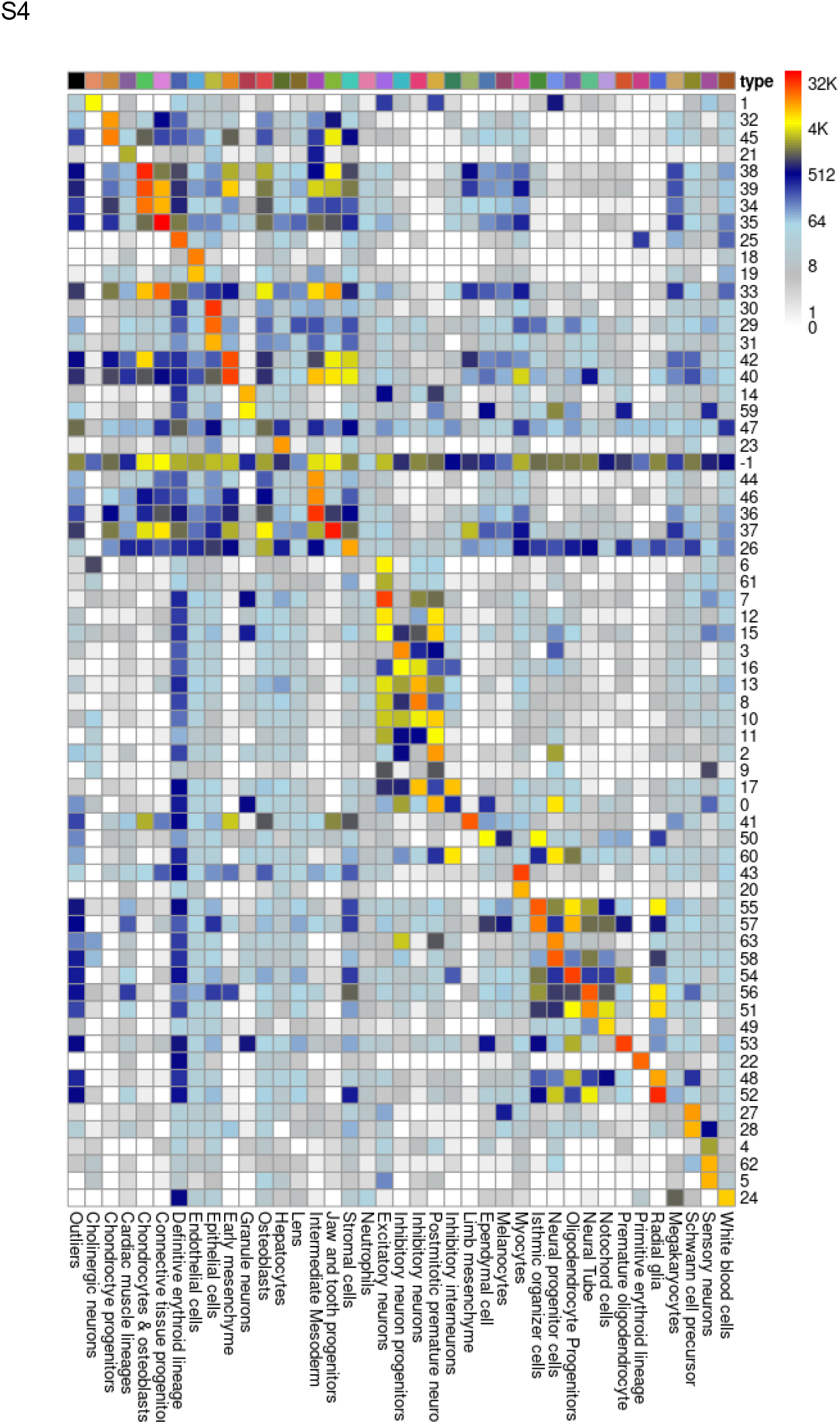
Organogenesis cell types vs. metacell clusters. Matrix is showing the number of cells in each combination of metacell cluster and organogenesis atlas cell type.

**Figure S5:**
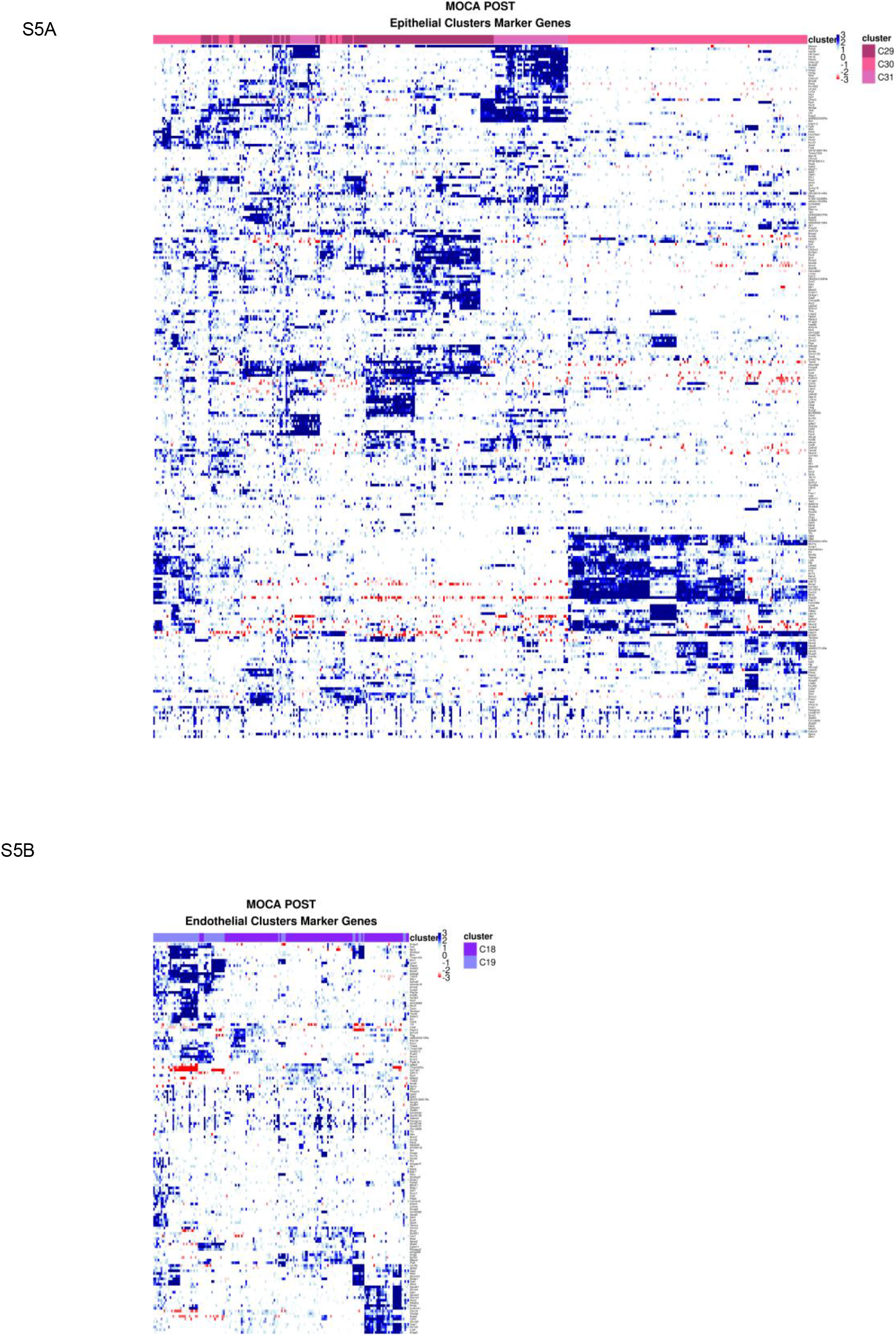
Metacell structure within broad cell types. Marker heat map (log2 gene expression normalized to the median over all metacell) is shown for metacells within the epithelial (29-31, A) and endothelial (18-19, B) metacell clusters. Rich combinatorial and quantitative variation is observed within each of the broader cell types, setting the stage for in-depth follow up analysis.

**Table S1.**

| Parameter | Default | Description |
| --- | --- | --- |
| rare_max_gene_cell_fraction | 0.001 (0.1%) | The maximal fraction of the cells where a gene is expressed to be considered "rare" |
| rare_min_gene_maximum | 7 | The minimal maximum-across-all-cells value of a gene to be considered as a candidate for rare gene modules |
| rare_min_genes_of_modules | 4 | The minimal number of genes in a rare gene module |
| rare_min_cells_of_modules | 12 | The minimal number of cells in a rare gene module |
| rare_min_cell_module_total | 4 | The minimal number of UMIs of a rare gene module in a cell to be considered as expressing the rare behavior |
| rare_max_cells_of_random_pile | 48 | The maximal mean number of cells in a random pile for a rare gene module to be considered rare |
| feature_downsample_min_samples | 750 | The minimal samples to use for downsampling the cells for computing "feature" genes |
| feature_downsample_min_cell_quantile | 0.05 (5%) | The minimal quantile of the cells total size to use for downsampling the cells for computing "feature" genes |
| feature_downsample_max_cell_quantile | 0.5 (50%) | The maximal quantile of the cells total size to use for downsampling the cells for computing "feature" genes |
| feature_min_gene_total | 50 | The minimal number of downsampled UMIs of a gene to be considered a "feature" |
| feature_min_gene_top3 | 4 | The minimal number of the top-3rd downsampled UMIs of a gene to be considered a "feature" |
| feature_min_gene_relative_variance | 0.1 | The minimal relative variance of a gene to be considered a "feature" |
| forbidden_gene_names | None | Genes forbidden from being a "feature" |
| forbidden_gene_patterns | None | Genes forbidden from being a "feature" |
| target_pile_size | 10000 | The target number of observations (cells) in a pile, allowing us to directly compute groups (metacells) for it |
| max_cell_size | None | The maximal cell size (total UMIs) to use |
| max_cell_size_factor | X 2 | The maximal cell size as a factor of the median cell size |
| cell_sizes | __x__\|sum<br>(Sum of UMIs) | The size of each cell for computing each metacell's size |
| target_metacell_size | 160000 | The target total metacell size (in UMIs) |
| knn_k | None<br>(Automatic) | The target K for building the K-Nearest-Neighbors graph |
| min_knn_k | 30 | The minimal target K for building the K-Nearest-Neighbors graph |
| candidates_cooldown_pass | 0.02 | By how much (as a fraction) to cooldown the temperature after doing a pass on all the nodes |
| candidates_cooldown_node | 0.25 | By how much (as a fraction) to cooldown the node temperature after improving it |
| candidates_cooldown_phase | 0.75 | By how much (as a fraction) to reduce the cooldown each time we re-optimize a slightly modified partition |
| candidates_min_metacell_cells | 12 | The minimal number of cells in a metacell, below which we would merge it |
| must_complete_cover | FALSE | Whether to force 100% coverage (disable outliers detection) |
| deviants_min_gene_fold_factor | 3<br>(X 8) | The minimal fold factor for a gene to indicate a cell is "deviant" |
| deviants_max_gene_fraction | 0.03<br>(3%) | The maximal fraction of genes to use to indicate cell are "deviants" |
| deviants_max_cell_fraction | 0.25<br>(25%) | The maximal fraction of cells to mark as "deviants" |
| dissolve_min_metacell_cells | 12 | The minimal number of cells in a metacell, below which we would dissolve it |
| random_seed | None<br>(Irreproducible) | Optional integer random seed for reproducible results |

## Notes

https://pypi.org/project/metacells/

### Competing Interest Statement

The authors have declared no competing interest.

## References

1. Picelli, S. et al. Full-length RNA-seq from single cells using Smart-seq2. Nat Protoc 9, 171–181 (2014).

2. Zheng, G. X. Y. et al. Massively parallel digital transcriptional profiling of single cells. Nat Commun 8, 14049 (2017).

3. Macosko, E. Z. et al. Highly Parallel Genome-wide Expression Profiling of Individual Cells Using Nanoliter Droplets. Cell 161, 1202–1214 (2015).

4. Svensson, V., Vento-Tormo, R. & Teichmann, S. A. Exponential scaling of single-cell RNA-seq in the past decade. Nat Protoc 13, 599–604 (2018).

5. Tanay, A. & Regev, A. Scaling single-cell genomics from phenomenology to mechanism. Nature 541, 331–338 (2017).

6. Kivioja, T. et al. Counting absolute numbers of molecules using unique molecular identifiers. Nat Meth 9, 72–74 (2012).

7. Jaitin, D. A. et al. Massively parallel single-cell RNA-seq for marker-free decomposition of tissues into cell types. Science (New York, N.Y.) 343, 776–9 (2014).

8. Pierson, E. & Yau, C. ZIFA: Dimensionality reduction for zero-inflated single-cell gene expression analysis. Genome Biology 16, 241 (2015).

9. Risso, D., Perraudeau, F., Gribkova, S., Dudoit, S. & Vert, J.-P. A general and flexible method for signal extraction from single-cell RNA-seq data. Nat Commun 9, 284 (2018).

10. Wagner, A., Regev, A. & Yosef, N. Revealing the vectors of cellular identity with singlecell genomics. Nat Biotechnol 34, 1145–1160 (2016).

11. Trapnell, C. et al. The dynamics and regulators of cell fate decisions are revealed by pseudotemporal ordering of single cells. Nat Biotechnol 32, 381–386 (2014).

12. Weinreb, C., Wolock, S., Tusi, B. K., Socolovsky, M. & Klein, A. M. Fundamental limits on dynamic inference from single-cell snapshots. PNAS 115, E2467–E2476 (2018).

13. Schiebinger, G. et al. Optimal-Transport Analysis of Single-Cell Gene Expression Identifies Developmental Trajectories in Reprogramming. Cell 176, 928-943.e22 (2019).

14. Haghverdi, L., Buettner, F. & Theis, F. J. Diffusion maps for high-dimensional singlecell analysis of differentiation data. Bioinformatics 31, 2989–2998 (2015).

15. Bergen, V., Lange, M., Peidli, S., Wolf, F. A. & Theis, F. J. Generalizing RNA velocity to transient cell states through dynamical modeling. Nat Biotechnol 38, 1408–1414 (2020).

16. Argelaguet, R. et al. MOFA+: a statistical framework for comprehensive integration of multi-modal single-cell data. Genome Biol 21, 111 (2020).

17. Setty, M. et al. Characterization of cell fate probabilities in single-cell data with Palantir. Nat Biotechnol 37, 451–460 (2019).

18. Gayoso, A. et al. Joint probabilistic modeling of single-cell multi-omic data with totalVI. Nat Methods 18, 272–282 (2021).

19. La Manno, G. et al. RNA velocity of single cells. Nature 560, 494–498 (2018).

20. Baran, Y. et al. MetaCell: analysis of single-cell RNA-seq data using K-nn graph partitions. Genome Biol. 20, 206 (2019).

21. Wolf, F. A., Angerer, P. & Theis, F. J. SCANPY: large-scale single-cell gene expression data analysis. Genome Biol 19, 15 (2018).

22. Stuart, T. et al. Comprehensive Integration of Single-Cell Data. Cell 177, 1888-1902.e21 (2019).

23. Fan, J. et al. Characterizing transcriptional heterogeneity through pathway and gene set overdispersion analysis. Nat Methods 13, 241–244 (2016).

24. Gayoso, A. et al. scvi-tools: a library for deep probabilistic analysis of single-cell omics data. http://biorxiv.org/lookup/doi/10.1101/2021.04.28.441833 (2021) xdoi:10.1101/2021.04.28.441833.

25. HCA Data Browser. https://data.humancellatlas.org/explore/projects/cc95ff89-2e68-4a08-a234-480eca21ce79?catalog=dcp1.

26. Cao, J. et al. The single-cell transcriptional landscape of mammalian organogenesis. Nature 566, 496–502 (2019).

27. Newman, M. E. J. & Girvan, M. Finding and evaluating community structure in networks. Phys. Rev. E 69, 026113 (2004).

28. Brandes, U. et al. On Modularity Clustering. (2008).

29. Blondel, V. D., Guillaume, J.-L., Lambiotte, R. & Lefebvre, E. Fast unfolding of communities in large networks. J. Stat. Mech. 2008, P10008 (2008).

30. Fogaça, M. et al. On the superiority of modularity-based clustering for determining placement-relevant clusters. Integration 74, 32–44 (2020).

31. Mouse RNA Atlas. https://oncoscape.v3.sttrcancer.org/atlas.gs.washington.edu.mouse.rna/downloads.

